# Reactive Astrocytes Prevent Maladaptive Plasticity after Ischemic Stroke

**DOI:** 10.1101/2021.06.16.448657

**Authors:** Markus Aswendt, Ulrika Wilhelmsson, Frederique Wieters, Anna Stokowska, Felix Johannes Schmitt, Niklas Pallast, Yolanda de Pablo, Lava Mohammed, Mathias Hoehn, Marcela Pekna, Milos Pekny

## Abstract

Restoration of functional connectivity is a major contributor to functional recovery after stroke. We investigated the role of reactive astrocytes in functional connectivity and recovery after photothrombotic stroke in mice with attenuated reactive gliosis (*GFAP^−/−^Vim^−/−^*). Infarct volume and longitudinal functional connectivity changes were determined by *in vivo* T2-weighted MRI and resting-state functional MRI. Sensorimotor function was assessed with behavioral tests, and glial and neural plasticity responses were quantified in the peri-infarct region. Four weeks after stroke, *GFAP^−/−^Vim^−/−^* mice showed impaired recovery of sensorimotor function and aberrant restoration of global neuronal connectivity. These mice also exhibited maladaptive plasticity responses, shown by higher number of lost and newly formed functional connections between primary and secondary targets of cortical stroke regions and increased peri-infarct expression of the axonal plasticity marker Gap43. We conclude that reactive astrocytes are required for optimal recovery-promoting plasticity responses after ischemic stroke.

## INTRODUCTION

Stroke is the second most common cause of death and the third leading cause of disability worldwide (Feigin et al., 2014; World Health Organization Foodborne Disease Burden Epidemiology Reference, 2012). The majority of stroke cases are due to cerebral artery occlusion, which triggers a pathophysiological cascade of cell death, neuroinflammation, and secondary neurodegeneration. Stroke-induced damage can lead to death or devastating functional deficit in the acute phase, but many stroke survivors achieve at least some functional recovery over months or years. Spontaneous functional improvement requires responses ranging from synaptic plasticity and axonal sprouting to re-organization of functional intra- and interhemispheric networks across the lesion border, leading to changes in existing neuronal pathways and new neuronal connections (Pekna et al., 2012). Stroke also triggers neurodegenerative changes in remote brain regions, such as the thalamus, that are not directly affected by the injury but are connected with the infarcted tissue. Clinical studies show that stroke-induced secondary neurodegeneration worsens the long-term outcome (Kuchcinski et al., 2017). Loss and gain of sensorimotor deficit–related functional connectivity and the spontaneous reorganization of functional neuronal networks can be monitored longitudinally by resting-state functional magnetic resonance imaging (rs-fMRI). *In vivo* rs-fMRI shows blood oxygen level–dependent signal fluctuations related to synchronized networks of spatially distinct brain regions at rest and is widely used to noninvasively monitor functional network changes after stroke in clinical studies (Thiel and Vahdat, 2015) and also in studies in rodents (Grandjean et al., 2020; Green et al., 2018; Mandino et al., 2019; Minassian et al., 2020). Reactive astrocytes are important cellular players in stroke (Li et al., 2008; Pekny and Nilsson, 2005; Pekny et al., 2019). In the acute phase, astrocytes protect brain tissue through mechanisms that support the restoration of brain homeostasis (Pekny et al., 2016; Pekny *et al.*, 2019; Verkhratsky and Nedergaard, 2018), and chemogenetic ablation of a subset of reactive astrocytes impairs vascular remodeling (Williamson et al., 2021). In the post-acute, functional recovery phase, reactive astrocytes modulate both axonal sprouting in the peri-infarct cortex and synaptic plasticity (Clarkson et al., 2010; Overman et al., 2012; Pekny *et al.*, 2019). However, the role of astrocytes in functional network changes after stroke is unknown.

Mice with genetic ablation of astrocyte intermediate filament (known also as nanofilament) proteins glial fibrillary acidic protein (GFAP) and vimentin (*GFAP^−/−^Vim^−/−^* mice) are a widely used model of attenuated reactive gliosis (Eliasson et al., 1999; Pekny et al., 1999b; Pekny and Pekna, 2014). In these mice, astrocytes do not become hypertrophic upon injury (Wilhelmsson et al., 2006; Wilhelmsson et al., 2004), and additional signs of attenuated reactive gliosis are present, such as reduced Erk or c-fos activation (Nakazawa et al., 2007) and reduced upregulation of 14-3-3 proteins (Sihlbom et al., 2007). Moreover, infarcts after middle cerebral artery occlusion are larger in *GFAP^−/−^Vim^−/−^* mice (Li *et al.*, 2008), and *GFAP^−/−^Vim^−/−^* astrocytes show decreased glutamine levels (Pekny et al., 1999a), lower glutamate transport (Li *et al.*, 2008), and impaired response to oxidative stress (de Pablo et al., 2013).

Here we investigated the role of reactive astrocytes in functional connectivity responses in *GFAP^−/−^Vim^−/−^* mice after ischemic stroke induced by photothrombosis. Sensorimotor function was assessed with behavioral tests, infarct volume and functional connectivity changes were determined by *in vivo* MRI, and glial and neural plasticity responses were quantified in the peri-infarct region.

## RESULTS

### Attenuation of reactive gliosis does not affect infarct size or acute loss of neuronal connectivity

*GFAP^−/−^Vim^−/−^* and wild-type (WT) mice were subjected to photothrombotic stroke in primary somatosensory and motor areas. Infarct size was quantified by *in vivo* T2-weighted MRI (T2w-MRI) and histology in combination with lesion mapping based on the Allen Brain Reference Atlas. In the acute and subacute phase, at post-stroke days 1 (P1) and 7 (P7), the ischemic lesion appeared as a hyperintense region on T2w-MRI. Lesion incidence maps showed low intragroup variability (**Figure 1B**). Lesion volume on T2w-MRI decreased between P1 and P7 (p < 0.001)); the groups did not differ in lesion volume (**Figure 1C**), shape (**Figures 1D and 1E**), or location (**Figures 1F and 1G**). At P1 in both *GFAP^−/−^Vim^−/−^* and WT mice, the lesion affected to a large extent (80 - 100%) the primary somatosensory cortex (SSp) and to a lesser extent (66 - 74%) the primary and secondary motor cortex (MOp and MOs). At 4 weeks, lesion volumes and regions affected were similar in the two groups (**Figures 1H–1J**). There was a strong correlation between *in vivo* and *ex vivo* measures of lesion size (**Table S1**).

**Figure 1.**
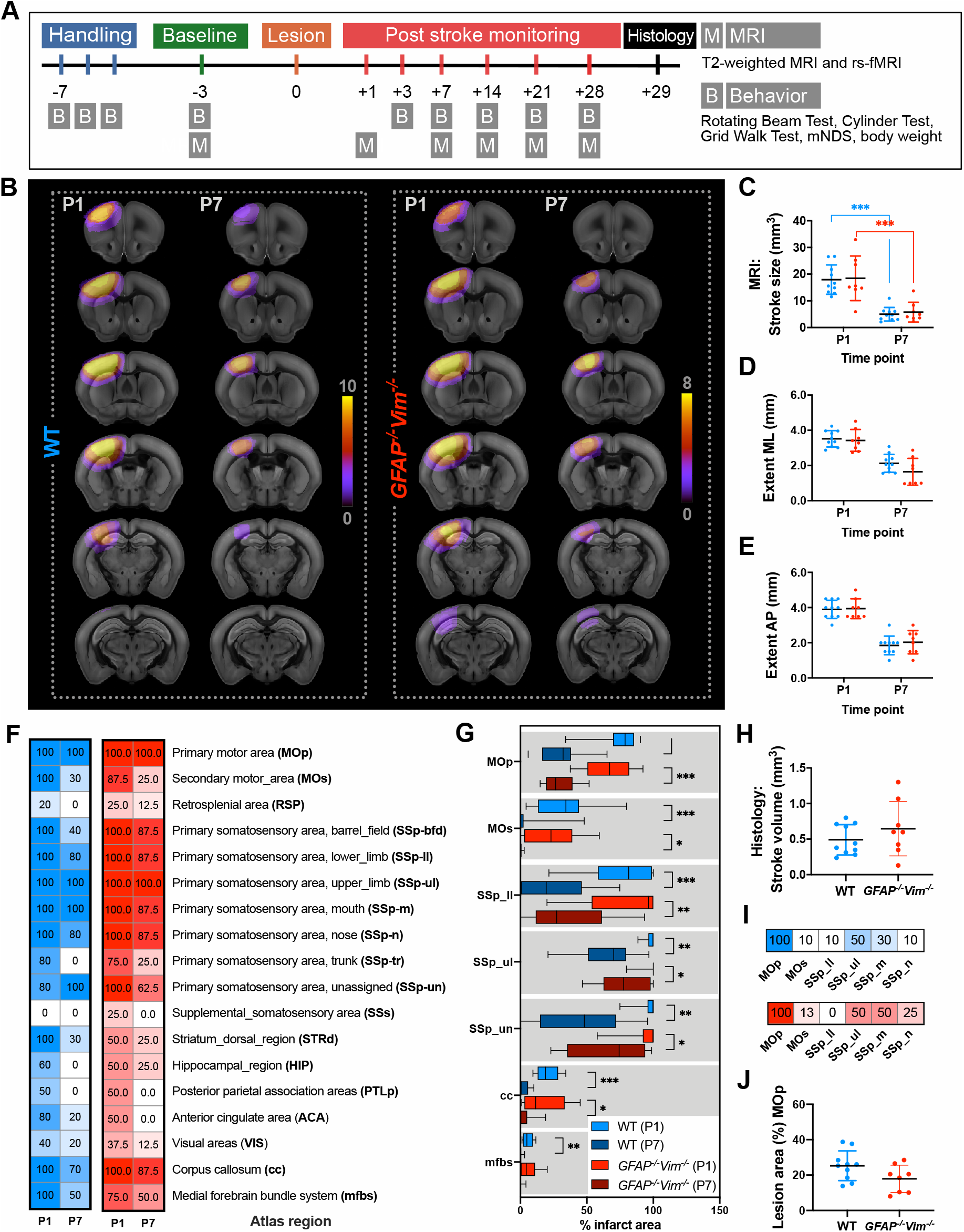
Attenuation of Reactive Gliosis in GFAP^−/−^Vim^−/−^ Mice Does Not Affect Infarct Size (A) Schematic of the study design. (B) MRI incidence maps: color indicates number of GFAP^−/−^Vim^−/−^ and WT mice in which a pixel was classified as part of a lesion area on T2w-MRI on P1 and P7. (C) Lesion volume determined by T2w-MRI. (D and E) Medial-lateral (ML) and anterior-posterior (AP) extent of the lesion determined on sagittal top-view projections of T2w-MRI. (F and G) Atlas-based mapping of T2w-MRI stroke lesions to individual brain regions. (F) incidence (% of mice affected) and (G) the extent of the region affected at P1 and P7. (H) Lesion volume determined from consecutive hematoxylin/eosin-stained brain sections at P29. (I and J) Atlas-based mapping of stroke lesions on hematoxylin/eosin-stained sections to individual brain regions with the incidence (% of mice affected) (I) and the extent of MOp affected (J) at P29. Data are presented as individual values with mean and SD (C–E, H, J), color map with percentage (F, I) or box plot (G). GFAP^−/−^Vim^−/−^ n=8, WT n=10. *p < 0.05, **p < 0.01, ***p < 0.001 by repeated-measures ANOVA with Sidak’s correction for multiple comparisons (C–F), a mixed-effects model with Tukey’s correction for multiple comparisons (L), and unpaired t test (H and J).

To investigate stroke-induced acute changes in the neuronal network response, we used atlas-based processing and graph theoretical analysis of rs-fMRI data. Acute consequences of the cortical lesions were evaluated according to the degree, representing the number of connections per node (brain region), and node strength, representing the sum of all edge weights (connectivity strength) per node. Globally (the entire brain network across 98 regions), connection degree and node strength (p < 0.001) decreased between baseline and P1, with no differences between groups (**Figures 2A and 2B**). The decrease connection degree and node strength was mostly restricted to sensorimotor-related regions (i.e. SSp-ll, STRv, and DORsm). Connectivity in the selected sensorimotor subnetwork (MOp, MOs, SSp-ul/ll/un), sensorimotor cortex–related thalamus (DORsm), polymodal association cortex–related thalamus (DORpm), dorsal striatum (STRd), and pallidum (PAL) was reduced at P1 (**Figures 2C–2F**). To assess the involvement of regions primarily affected by stroke, we compared connection degree and node strength in SSp-ul/ll/un and MOp/s (**Figures 2G and 2H**). Both measures decreased from baseline to P1 in WT mice (connection degree: p < 0.001, node strength: p < 0.001) and *GFAP^−/−^Vim^−/−^* mice (connection degree: p < 0.01, node strength: p < 0.001), with no differences between groups at baseline or P1.

**Figure 2.**
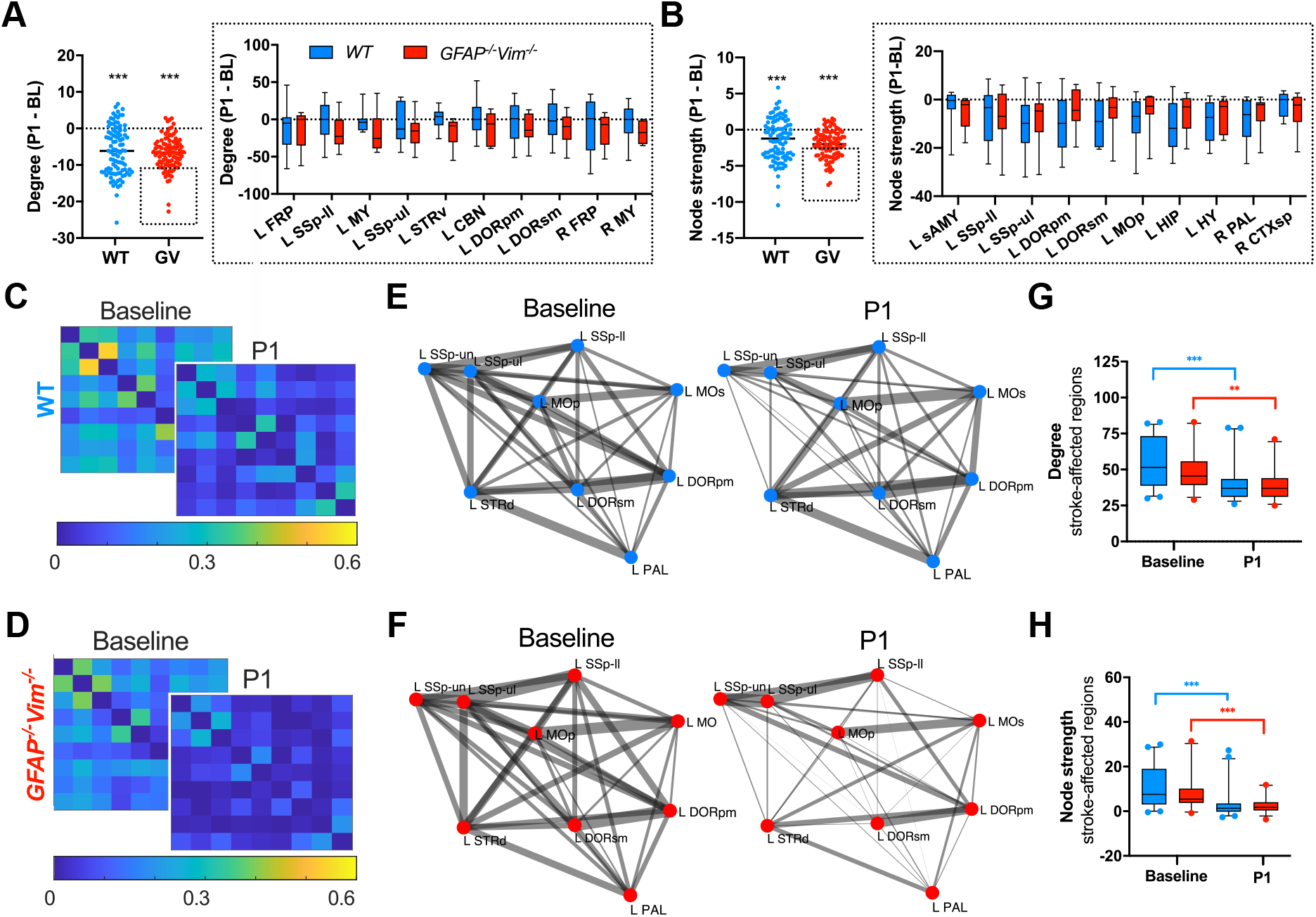
Attenuation of Reactive Gliosis Does Not Affect the Acute Loss of Neuronal **Connectivity** Acute stroke-induced neuronal network changes were determined by rs-fMRI in WT (blue) and GFAP^−/−^Vim^−/−^ (GV, red) mice. (A) Total degree (number of network connections in all brain regions) on P1 normalized to baseline (BL) and regions with the largest change in connection degree (top 10%). (B) Total node strength (the strength of connectivity in all brain regions) at P1 normalized to baseline and regions with the largest change in node strength (top 10%). (C–F) Sensorimotor subnetwork (brain regions SSp-ll/ul/un, MOs/p, PAL, STRd, DORsm/pm) connectivity at baseline and P1 visualized by mean correlation matrices (C and D) for WT (C) and GFAP^−/−^Vim^−/−^ (D) mice and by graphs of quantitative data for the same network (E and F). Circles indicate nodes; line thickness indicates edge weights. (G and H) Quantitative analysis of connection degree and node strength in the areas primarily affected by stroke (SSp-ul/ll/un and MOp/s) at baseline and P1. Data are presented as individual values (A, B) or box plots (G, H). n = 7 GFAP^−/−^Vim^−/−^ mice; n = 9 WT mice. **p < 0.01, ***p < 0.001 by paired t test (baseline vs P1), unpaired t test (between groups), two-way ANOVA with FDR correction for region-wise comparisons (A, B), and mixed-effects model with FDR correction for between time points or intergroup comparisons (G, H).

### Attenuation of reactive gliosis impairs functional recovery after ischemic stroke

To assess stroke-induced impairment and spontaneous functional recovery, we used the rotating beam, cylinder, and grid walk tests to evaluate sensorimotor function at baseline and for 4 weeks after stroke. Functional impairment relative to baseline was expressed as number of hindlimb drops and frequency of paw drags and foot faults. Starting on P7, the *GFAP^−/−^Vim^−/−^* mice showed more hindlimb drops, paw drags, and foot faults than WT mice (p < 0.05, p < 0.01, and p < 0.001, respectively, on P28) (**Figures 3A–3C**). In WT mice, hindlimb drops and paw drags quickly decreased and did not differ from pre-stroke levels after P7. In contrast, in *GFAP^−/−^Vim^−/−^* mice, the number of hindlimb drops and paw drags remained increased on P21 (p < 0.05) and P28 (p < 0.05), respectively. The frequency of foot faults in the grid walk test at P28 was still higher than pre-stroke levels in both groups (*GFAP^−/−^Vim^−/−^*: p < 0.001, WT: p < 0.01). However, WT mice improved between P14 and P28 (p < 0.001), whereas *GFAP^−/−^Vim^−/−^* mice did not (**Figure 3C**). The combined average sensorimotor deficit calculated from all three measures on P3 was similar in the two groups, but the rate of improvement was lower in *GFAP^−/−^Vim^−/−^* mice (R^2^ = 0.710/slope = –1.858 vs. R^2^ = 0.697/slope = –2.551; p < 0.05) (**Figure 3D**). All behavioral measures were independent of lesion volume determined by T2-weighted MRI on P1 and by histology on P28 (**Table S1**).

**Figure 3.**
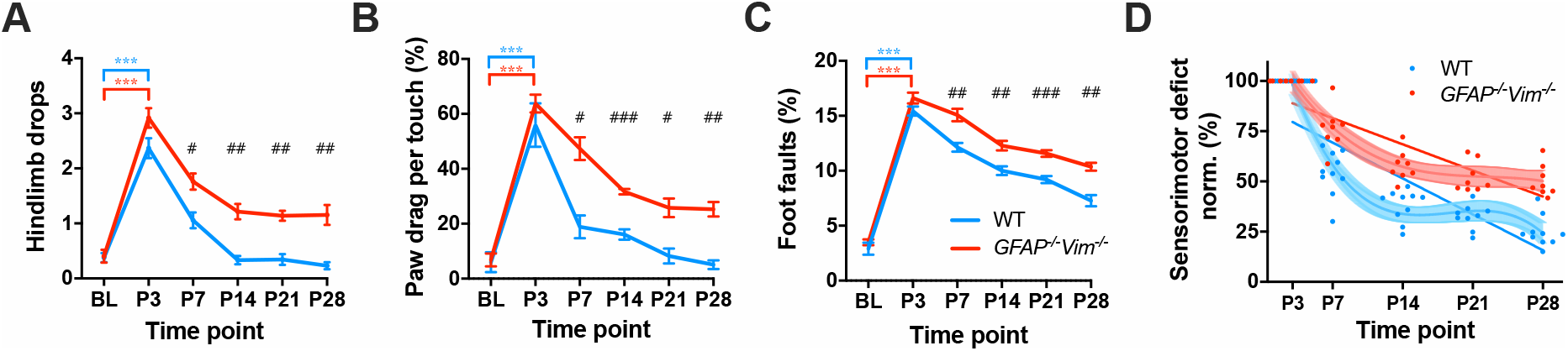
Attenuation of Reactive Gliosis Impairs Functional Recovery after Ischemic Stroke. Evaluation of post-stroke sensorimotor function. (A–C) Functional recovery as assessed by the number of hindlimb drops in the rotating beam test, relative frequency of paw drags in the cylinder test, and the fraction of foot faults in the grid walk test. (D) Combined average functional deficit based on rotating beam hindlimb drop, cylinder test paw drag, and grid walk foot fault data for all WT and GFAP^−/−^Vim^−/−^ mice. Red and blue lines represent linear regression with a significantly different slope between the groups (R^2^WT = 0.697 and R^2^GV = 0.710). Light red and light blue segments represent nonlinear, third-order polynomial fit including the 95% confidence level (R^2^WT = 0.923 and R^2^GV = 0.874). Data are presented as mean ± SEM (A–C) or individual values (D). Rotating beam: GFAP^−/−^Vim^−/−^ n = 7, WT n = 10. Grid walk: GFAP^−/−^Vim^−/−^ n = 8, WT n = 10. Cylinder test: GFAP^−/−^Vim^−/−^ n = 6, WT n = 10. WT, blue lines; GFAP^−/−^Vim^−/−^, red lines. Mixed-effects model with Dunnett’s and Sidak’s correction for between time points (***p < 0.001) and intergroup comparisons (^#^p < 0.05, ^##^p < 0.01, and ^###^p < 0.001), respectively.

### Attenuation of reactive gliosis alters stroke-induced neuronal connectivity responses in the post-acute phase

To assess post-stroke functional connectivity changes, we analyzed rs-fMRI data from the late post-stroke stage. At P28, connection degree across the whole brain was lower **(Figure 4A)** than at baseline in *GFAP^−/−^Vim^−/−^* mice (p < 0.01); the median was –3.5. In contrast, connection degree in WT mice reached baseline levels by P28 (p < 0.001 vs *GFAP^−/−^Vim^−/−^*). Analysis of the regions in which the decrease in connection degree relative to baseline was in the top 10% in *GFAP^−/−^Vim^−/−^* mice showed no overall difference between *GFAP^−/−^Vim^−/−^* and WT mice. Both groups had lower node strength at P28 than at baseline (p < 0.001), but the decrease was larger in *GFAP^−/−^Vim^−/−^* mice (median –4.3 vs. –2.2, p < 0.001) **(Figure 4B)**.

**Figure 4.**
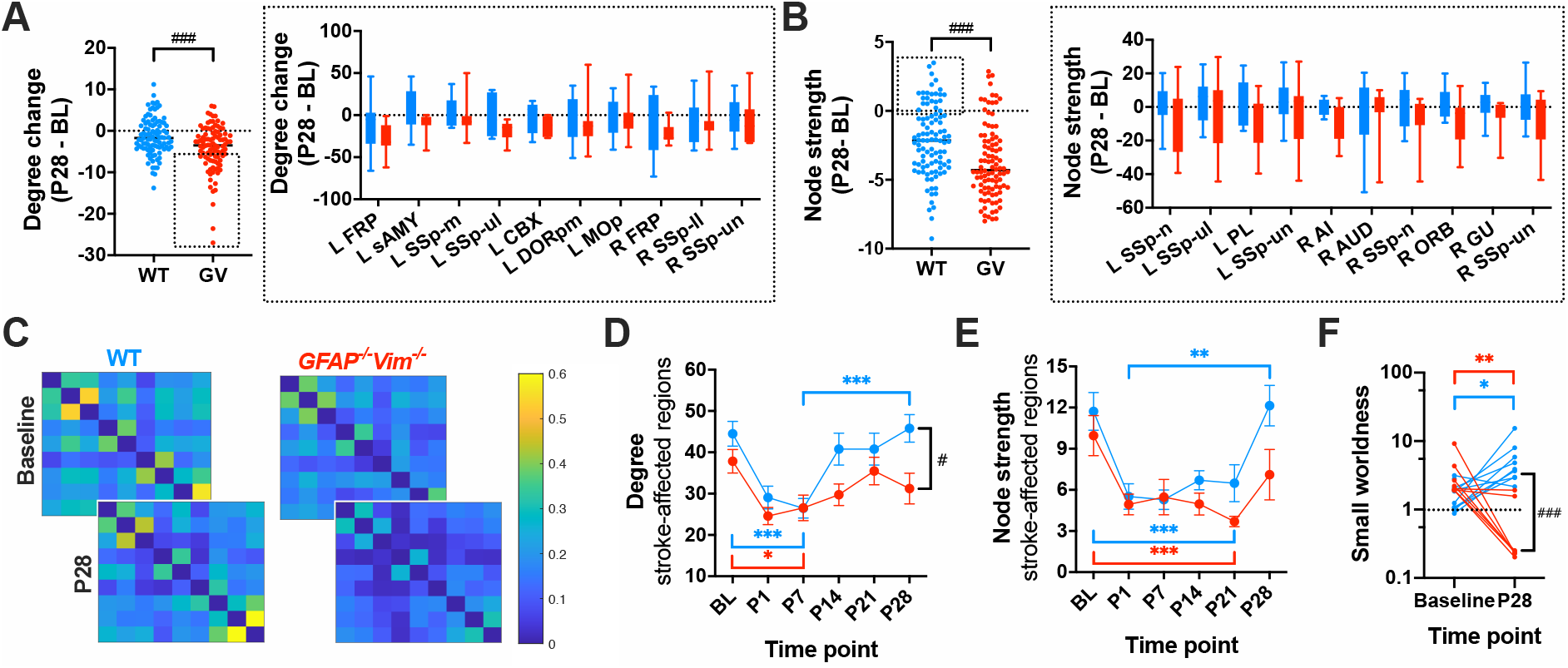
Attenuation of Reactive Gliosis Alters Stroke-Induced Neuronal Connectivity **Responses in the Post-acute Phase** Late post-stroke neuronal network changes determined by rs-fMRI in WT (blue) and GFAP^−/−^Vim^−/−^ (GV, red) mice. (A) Change in total degree (the number of network connections in all brain regions) at P28 relative to baseline in GFAP^−/−^Vim^−/−^ and WT mice, with box and bar graph plot for the top 10% of negative regions in GFAP^−/−^Vim^−/−^ mice and the respective regions in WT mice. (B) Change of total node strength (the strength of connectivity in all brain regions) at P28 relative to baseline for GFAP^−/−^Vim^−/−^ and WT mice, with box and bar graph plot for the top 10% of positive regions in GFAP^−/−^Vim^−/−^ mice and the respective regions in WT mice. (C) Mean correlation matrices for GFAP^−/−^Vim^−/−^ and WT mice at baseline and P28 in selected sensorimotor regions (SSp-ul/ll/un, MOp/s, STRd, PAL, DORsm). (D and E) Longitudinal change in connection degree (D) and node strength (E) in the regions primarily affected by stroke (SSp-ul/ll/un, MOp/s, and DORpm/sm). (F) Change in small-worldness from baseline to P28. GFAP^−/−^Vim^−/−^ n = 7, WT n = 9. *p < 0.05, **p < 0.01, ***p < 0.001 by paired t test (baseline vs P28), unpaired t test (between groups), two-way ANOVA with false-discovery rate correction for region-wise comparisons (A, B), and mixed-effects model with false-discovery rate correction for time point comparisons or intergroup comparisons (D–F).

Next, we performed region-specific analysis of rs-fMRI data from the sensorimotor network and the regions primarily affected by stroke (**Figures 4C–4E**). In both groups, the average connection degree in affected regions (MOp, MOs, SSp-ul/ll/un, DORsm, DORpm, STRd and PAL) decreased until P7 (WT: p < 0.001, *GFAP^−/−^Vim^−/−^*: p < 0.05) but did not differ from baseline on P14, P21, or P28. In the WT group, however, connection degree increased from P7 to P14 (p < 0.01) and continued to increase until P28 (p < 0.001), when it was higher than in *GFAP^−/−^Vim^−/−^* mice (p < 0.05). In both groups, average node strength in stroke-affected regions at P7, P14, and P21 was lower than at baseline (p < 0.001 at P21) and reached baseline levels at P28 with no intergroup differences. However, only in WT mice did node strength increase between P1 and P28 (p < 0.01).

Network small-worldness, a measure of local connectivity, was similar in the two groups at baseline (**Figure 4F)**. At P28, however, it had increased in WT mice (p < 0.01) but decreased in *GFAP^−/−^Vim^−/−^* mice (p < 0.05) and was lower than in WT mice (p < 0.001), suggesting impaired connectivity between neighboring brain regions. In the acute phase, network changes in stroke-affected regions did not correlate with sensorimotor function; at P28, however, connection degree, node strength, and small-worldness correlated negatively with the number of hindlimb drops and frequency of paw drags and foot faults **(Table 1)**.

**Table 1.** Results of Spearman correlation (Spearman r_s_ and p-value) of connection degree, node strength, and small-worldness in stroke-affected regions (MOp/s, SSp-ul/ll/un) with the functional readouts hindlimb drop (rotating beam test), paw drag (cylinder test), and foot fault (grid walk test).

|  | **P1 degree**<br>**vs. P3 hindlimb drop** | **P1 degree**<br>**vs. P3 paw drag** | **P1 degree**<br>**vs. P3 foot fault** |
| --- | --- | --- | --- |
| $r_s$ | 0.261 | 0.106 | 0.205 |
| p | 0.327 | 0.683 | 0.414 |
|  | **P1 node strength**<br>**vs. P3 hindlimb drop** | **P1 node strength**<br>**vs. P3 paw drag** | **P1 node strength**<br>**vs. P3 foot fault** |
| $r_s$ | -0.224 | 0.349 | -0.345 |
| p | 0.480 | 0.219 | 0.227 |
|  | **P28: degree**<br>**vs. hindlimb drop** | **P28: degree**<br>**vs. paw drag** | **P28: degree**<br>**vs. foot fault** |
| $r_s$ | -0.6324 | -0.5608 | -0.325 |
| p | 0.008** | 0.026* | 0.188 |
|  | **P28: Node strength**<br>**vs. hindlimb drop** | **P28: Node strength**<br>**vs. paw drag** | **P28: Node strength**<br>**vs. foot fault** |
| $r_s$ | -0.242 | -0.570 | -0.679 |
| p | 0.402 | 0.036* | 0.007** |
|  | **P28: Small-worldness**<br>**vs. hindlimb drop** | **P28: Small-worldness**<br>**vs. paw drag** | **P28: Small-worldness**<br>**vs. foot fault** |
| $r_s$ | -0.760 | -0.501 | -0.566 |
| p | 0.001*** | 0.059 | 0.020* |

### Stroke alters specific connections in the sensorimotor network in WT mice

To determine whether the functional neuronal network was re-established with all individual connections preserved or whether individual connections were altered or even replaced, we used rs-fMRI network analysis as a novel approach to assess the individual connections of stroke-affected brain regions. We focused on the 21 strongest connections (highest inter-node correlation strength at baseline) of the sensorimotor networks (SSp-ul/ll/un, SSs, DORsm/pm, MOp/s, STRd) in the ipsilesional (left) and contralesional (right) hemispheres (**Table S2**). In the WT group, 64.0 ± 9.3% of the strongest intra- and interhemispheric connections were preserved or re-established **(Figure 5).** Specifically, in SSp-ll (directly affected by stroke) and STRd (secondarily affected), four baseline connections were lost, whereas five were new at P28. Thus, in WT mice, specific connections were lost after stroke, and new ones were established. This approach provides more specific information about stroke-induced sensorimotor network responses.

**Figure 5.**
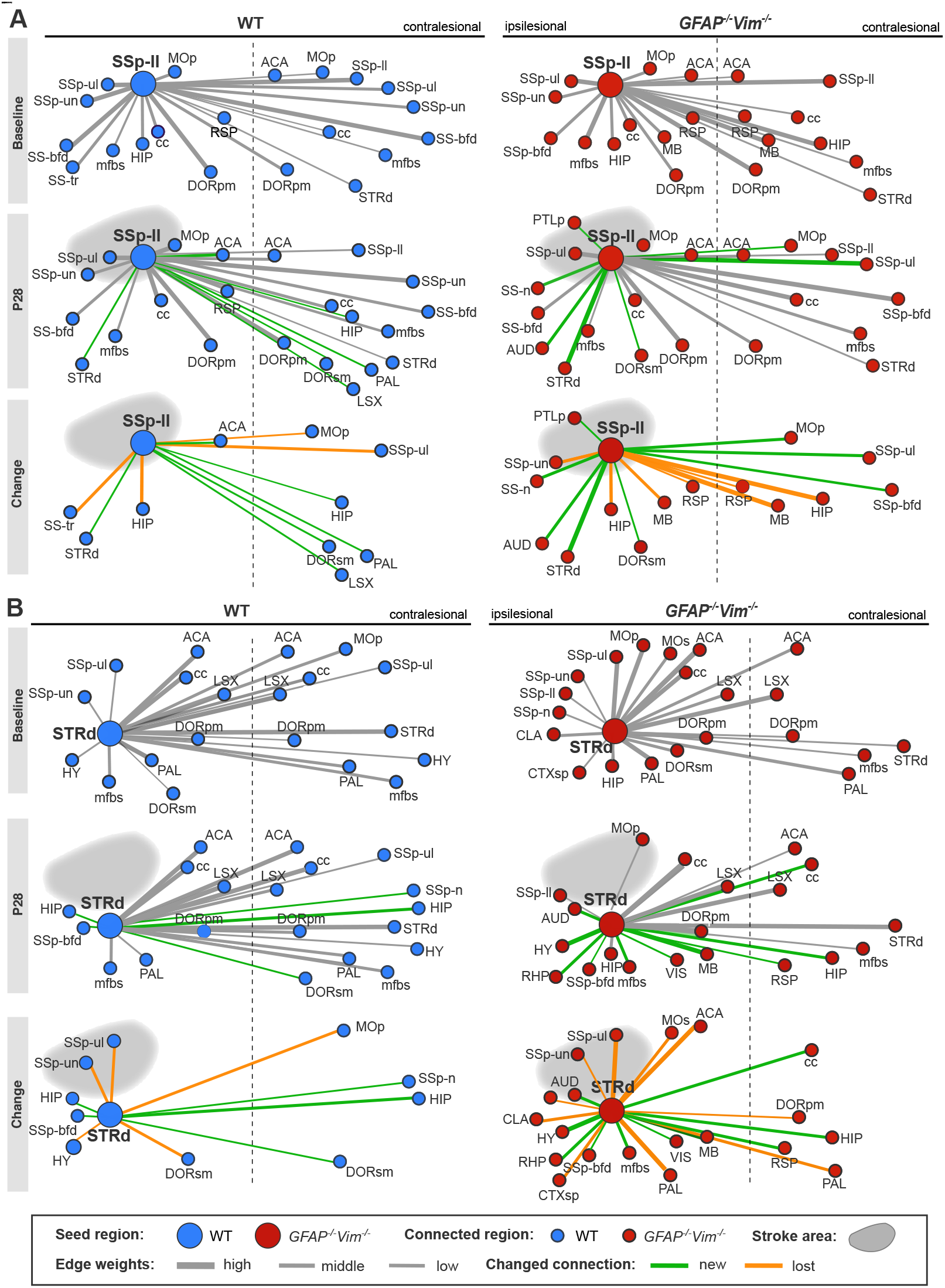
Reactive Gliosis-Dependent Change in Functional Connectivity after Stroke in **Primary and Secondary Targets of Cortical Stroke** (A and B) Illustration of the top 21 connections of ipsilesional SSp-ll (A) and ipsilesional STRd (B). Thresholding was based on the group average z-score correlations of rs-fMRI data at baseline (top) and on P28 (middle); the bottom part highlights newly formed and lost connections at P28. Line thickness indicates connection strength (high, middle, and low) (extended list in Table S2).

### Attenuation of reactive gliosis increases loss and gain of connections in sensorimotor networks

Next, we applied the same approach to sensorimotor networks in *GFAP^−/−^Vim^−/−^* mice. At P28, fewer of the strongest intra- and interhemispheric connections were preserved or re-established than in WT mice (49.5 ± 8.7% vs 64.0 ± 9.3%) **(Figure 5)**. Specifically, in SSp-ll, *GFAP^−/−^Vim^−/−^* mice lost more connections (7 vs 4) and had more new connections (8 vs 5). Similarly, in STRd, *GFAP^−/−^Vim^−/−^* mice lost more connection (10 vs 5) and had more new connections (10 vs 5). At P28, only 7.7 ± 9.0% of the new strongest connections (green in Figure 5) were identical in both groups. Importantly, in *GFAP^−/−^Vim^−/−^* mice, the number of connections that did not change between baseline and P28 was lower in both the ipsilesional (p < 0.01) and contralesional (p < 0.001) networks. Thus, attenuation of reactive gliosis impaired the restoration of normal sensorimotor networks and induced new connections different from those in WT mice.

### Impaired polarization of astrocyte processes and decreased peri-lesion astrocyte density in GFAP^−/−^Vim^−/−^ mice

To assess the post-stroke response of glial cells, we used immunohistochemistry to identify and quantify Iba1^+^ microglia and Sox2^+^ or S100^+^ astrocytes in the cortex at P29 (**Figure 6**). The densities of Iba1^+^ microglia and Sox2^+^ astrocytes were similar in *GFAP^−/−^Vim^−/−^* and WT mice, and in both groups the density of Iba^+^ cells was higher in the ipsilesional than contralesional somatosensory cortex (**Figures 6A–6C**). Immunostaining for S100 suggested reduced polarity of astrocytes in response to the lesion in *GFAP^−/−^Vim^−/−^* mice (**Figure 6D**). In WT mice, the directionality of S100^+^ astrocyte processes showed polarization of astrocytes toward the lesion in the ipsilesional cortex. This polarization response was impaired in *GFAP^−/−^Vim^−/−^* mice (p < 0.05) (**Figures 6E and 6F**), and S100^+^ profiles in ipsilesional cortex were shorter than in WT mice (**Figure 6G**). In the peri-infarct cortex adjacent to infarcted tissue (**Figure 6H**), microglia activation was similar in the two groups, as judged from the Iba1^+^ area (**Figure 6I**), whereas Sox2^+^ astrocytes in the same area were less abundant in *GFAP^−/−^Vim^−/−^*mice (**Figure 6J**). Thus, astrocyte density at the edge of the lesion is decreased and the polarization of astrocyte processes is impaired in *GFAP^−/−^Vim^−/−^* mice, confirming attenuated reactive gliosis. The overall cortical densities of microglia and astrocytes at P29 were not affected by attenuation of reactive gliosis.

**Figure 6.**
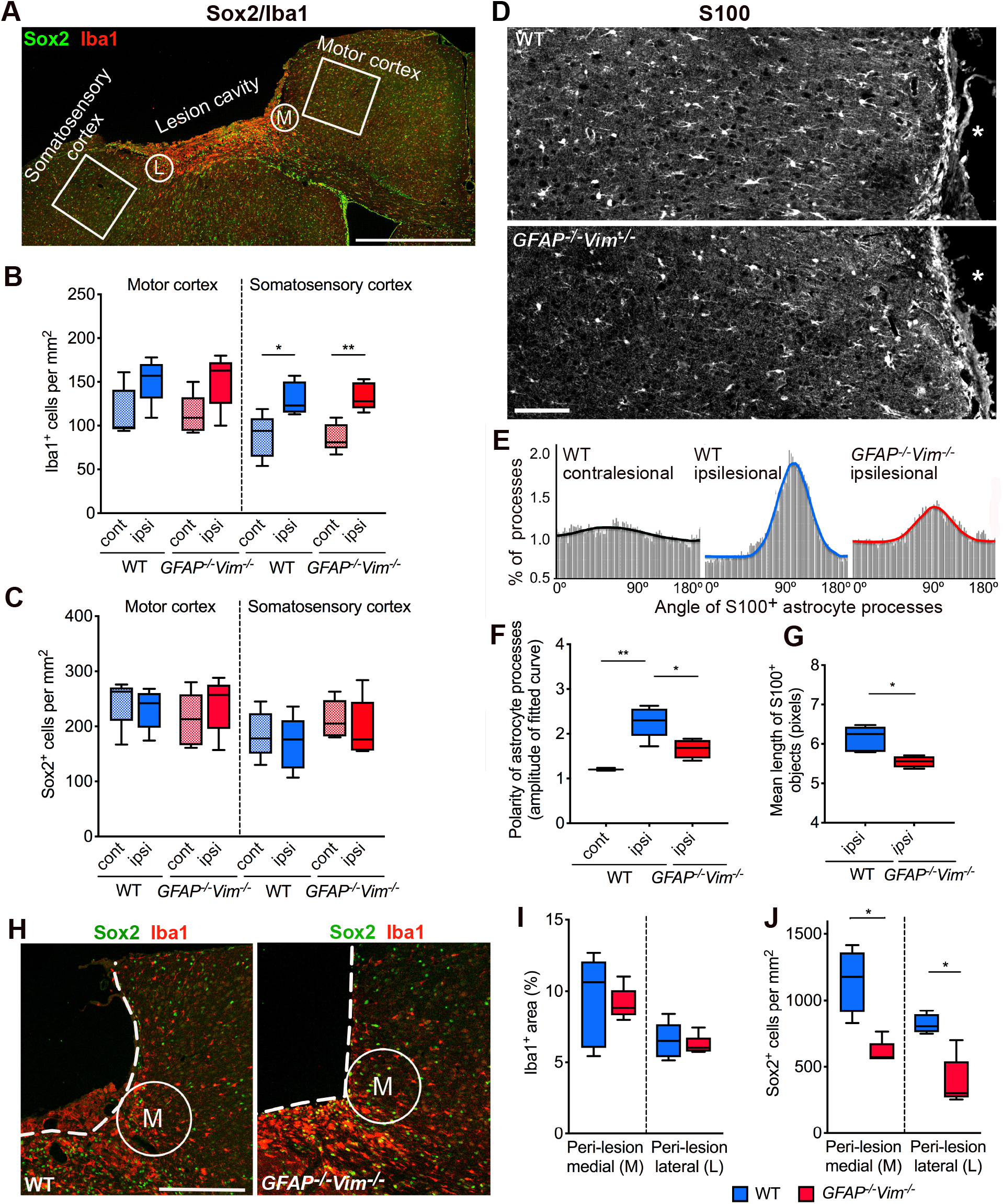
Impaired Polarization of Astrocyte Processes and Decreased Peri-Lesion Astrocyte **Density in GFAP^−/−^Vim^−/−^ Mice** (A) Coronal section of ipsilesional cortex 4 weeks after stroke. Green, Sox2^+^ astrocytes; red, Iba1^+^ microglia. Squares and circles indicate areas used for quantification. (B and C) Density of Iba1^+^ cells (B) and Sox2^+^ cells (C) in the squares within ipsilesional (ipsi) and contralesional (cont) cortex. (D) Coronal section of peri-lesional cortex 4 weeks after stroke; astrocytes and their processes were visualized by S100 immunohistochemistry. Asterisk denotes lesion cavity. (E–G) Polarity of astrocyte processes. Representative histograms show process directionality (E), amplitude of the fitted curve (F), and length of thresholded S100^+^ objects (G). **(H and J)** Intensity of Iba1^+^ immunostaining (I) and Sox2^+^ cell density (J) in the medial (M) and lateral (L) peri-lesional cortex in the area denoted by circles in A and D. Dotted line demarcates the lesion. Scale bars: A, 1000 μm; D, 100 μm; H, 250 μm. n = 5 (B, C, I, J) or 3–5 (F, G) per group. Data are presented as box plots. *p < 0.05, **p < 0.01 by one-way ANOVA with Sidak’s or Tukey’s correction for multiple comparisons (B, C, F, G) and Mann-Whitney test (I, J).

### Attenuation of reactive gliosis alters neural plasticity responses in the peri-infarct region

To assess stroke-induced plasticity changes on a tissue level at 4 weeks, we identified presynaptic terminals of excitatory neurons with antibodies against glutamate transporter 1 (Vglut1) (**Figure 7A**). In WT mice, high-content image analysis showed increases in the number, size, and signal intensity of Vglut1-positive puncta in cortical layers V-VI of the motor and somatosensory peri-lesional cortex vs contralesional cortex (**Figures 7B–7D**). In *GFAP^−/−^Vim^−/−^* mice, however, the only difference between these areas was the greater size of Vglut1^+^ puncta in the ipsilesional motor cortex, indicating a weaker stroke-induced synaptogenic response (**Figures 7B–7D**). Finally, we used immunohistochemistry to assess expression of Gap43, a marker of axonal and glial plasticity (**Figure 7E**). The density, size, and intensity of Gap43^+^ puncta in layers V-VI of the peri-infarct motor cortex were greater in *GFAP^−/−^Vim^−/−^* compared to WT mice (p < 0.05, p < 0.05, and p < 0.001, respectively), indicating higher expression of Gap43 (**Figures 7F–7H**). The density and intensity of the Gap43^+^ puncta in layers V-VI was also higher in the peri-infarct somatosensory cortex of *GFAP^−/−^Vim^−/−^* compared to WT mice (both p < 0.05). These results suggest that glial and axonal plasticity responses are increased by attenuation of reactive gliosis.

**Figure 7.**
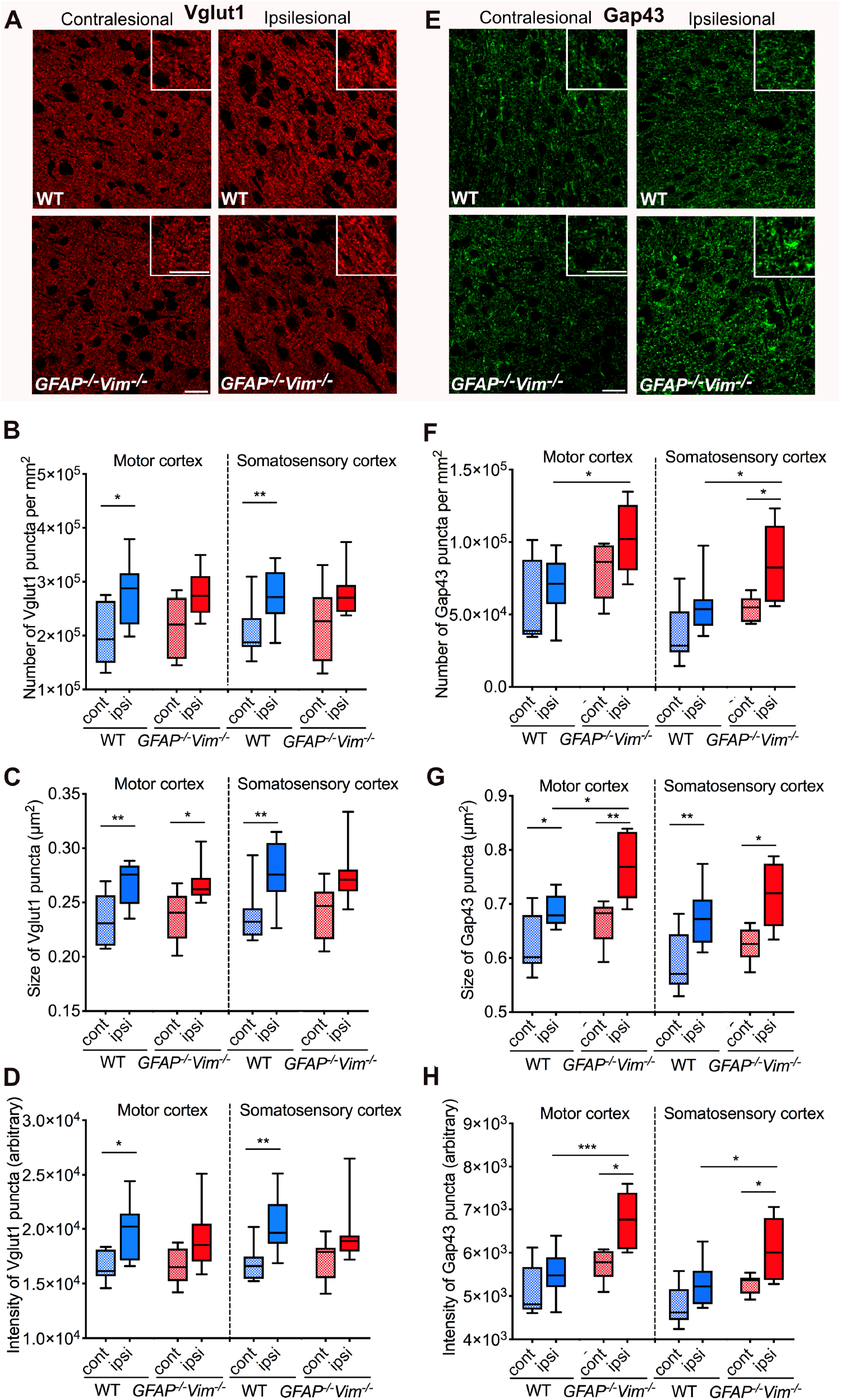
Attenuation of Reactive Gliosis Alters Neural Plasticity Responses in the Peri-infarct **Cortex.** (A–H) Representative images of Vglut1 (A) and Gap43 (E) immunoreactivity in the ipsilesional and contralesional cortex at 4 weeks after stroke; insets show high-magnification images. Quantification of Vglut1 (B–D) and Gap43 (F–H) immunoreactivity as puncta density (upper), size (middle), and intensity (lower). Scale bars A and E, 20 μm. Cont, contralesional; ipsi, ipsilesional. A–D, n = 8–9 per group; E–H, n = 6–9 per group. Data are presented as box plots. *p < 0.05, **p < 0.01, ***p < 0.001 by one-way ANOVA with Sidak’s correction for multiple comparisons.

## DISCUSSION

Restoration of functional connectivity is important for functional recovery after stroke (Green *et al.*, 2018; Pan et al., 2015; van der Zijden et al., 2008a; van der Zijden et al., 2008b; van Meer et al., 2010), but the mechanisms of functional brain re-organization after stroke are only partially understood. This study shows that attenuation of reactive gliosis leads to an aberrant reorganization of both the global and the affected sensorimotor functional network after cortical stroke. These results suggest that reactive gliosis is necessary for normalization of functional neuronal connectivity and for plasticity responses in the peri-infarct cortex that promote functional recovery. We found that attenuated reactive gliosis does not affect lesion size or the initial functional deficit after stroke. As in previous rs-fMRI studies of stroke-induced changes in functional connectivity (Carter et al., 2010; Golestani et al., 2013; van Meer *et al.*, 2010), functional connectivity in the acute phase after stroke was strongly reduced. This acute reduction in functional connectivity was similar in *GFAP^−/−^Vim^−/−^* and WT mice. During the recovery phase, the degree and node strength of functional connectivity did not reach pre-stroke levels in mice with attenuated reactive gliosis, and their behavioral recovery was slower and less complete. These findings suggest a less stable functional network characterized by more and stronger stroke-induced new connections and by greater stroke-induced loss of connectivity. Moreover, functional outcome correlated positively with both the connection degree and node strength of the entire brain network, consistent with the correlation between normalized activation patterns in fMRI and functional recovery in chronic stroke patients (Ward et al., 2003). These findings further implicate restoration of neuronal connectivity in post-stroke recovery of function and the critical role of reactive astrocytes in that process.

Although WT mice achieved a functional normalization of the neuronal network on the whole-brain level, our detailed sensorimotor network topology analysis revealed that stroke causes long-lasting alterations in the number and strength of connections in the primarily affected brain regions and other regions of the sensorimotor network. The massive loss of originally strong connections and the strengthening of weak connections—many from stroke-affected regions to contralesional, nonsensorimotor regions—in mice with attenuated reactive gliosis point to a suboptimal or maladaptive gain and loss of top connections after stroke. Our results also point to a major role of reactive gliosis in restoring network small-worldness, a measure of high clustering and short path length within a network, indicative of fast propagation of information and strong synchronizability between network nodes (Watts and Strogatz, 1998). The differences in small-worldness between *GFAP^−/−^Vim^−/−^* and WT mice and the negative correlation of small-worldness with the extent of impairment 4 weeks after stroke further support the notion that reactive gliosis is required for optimal network re-organization after stroke.

There is a growing list of molecules whose expression is induced in the peri-infarct region and which have been implicated as positive and negative regulators of post-stroke connectivity changes (Murphy and Corbett, 2009). Many of these factors and their receptors, such as thrombospondins, ephrin A5, and semaphorin 3A, are expressed by reactive astrocytes (Pekny and Pekna, 2014; Pekny *et al.*, 2019). We and others have shown that reactive gliosis limits a range of neuroplasticity and regenerative responses in the CNS, such as synaptic plasticity (Wilhelmsson *et al.*, 2004), axonal regeneration (Cho et al., 2005; Menet et al., 2003), baseline or pathology-triggered neurogenesis (Jarlestedt et al., 2010; Wilhelmsson et al., 2012), and integration of neural grafts (Kinouchi et al., 2003; Widestrand et al., 2007).

Synaptic plasticity and functional remapping in the peri-infarct cortex and in the contralesional hemisphere were proposed as important factors in post-stroke functional recovery (reviewed in Pekna *et al.*, 2012). Post-stroke synaptic plasticity in the peri-infarct cortex is associated with the reactivation of an intrinsic neuronal growth program and upregulation of Gap43 (Carmichael et al., 2005), a membrane phosphoprotein associated with axonal growth cones and a marker of axonal sprouting (Benowitz et al., 1990; Benowitz and Routtenberg, 1997). Gap43 in the peri-infarct cortex is predominantly localized in the neuronal compartment, near pre-synaptic terminals, and in astrocytes (Stokowska et al., 2017). Gap43 expressed in astrocytes mediates glial plasticity during reactive gliosis, attenuates microglial activation, and fosters neuronal survival and plasticity (Hung et al., 2016). Our findings— specifically, the altered expression of Gap43 and the weaker synaptogenic response seen as the relative increase in the density of Vglut1^+^ presynaptic terminals of excitatory neurons in the peri-infarct region of mice with attenuated reactive gliosis—further support the notion that reactive astrocytes help modulate post-stroke neural plasticity. This altered synaptogenic response after stroke in mice with attenuated reactive gliosis is in line with the more pronounced deafferentation injury–induced loss and faster normalization of the number of synapses in the hippocampus of these mice (Wilhelmsson *et al.*, 2006; Wilhelmsson *et al.*, 2004). Given the impaired recovery of sensorimotor function in mice with attenuated reactive gliosis, these neural plasticity responses appear to be maladaptive, leading to suboptimal or random neuronal network reorganization. As random spontaneous axonal outgrowth was proposed as the mechanism of less optimal network organization (Thiel and Vahdat, 2015), the increased expression of Gap43 and the reduced astrocyte polarity in the peri-infarct cortex of mice with attenuated reactive gliosis implicate reactive gliosis as an inhibitor of maladaptive plasticity responses in the post-acute phase after stroke.

Our results point to a novel role of reactive gliosis in restoration of functional connectivity and recovery after stroke. By virtue of their neuroprotective functions in the acute phase and their ability to promote adaptive plasticity in the post-acute phase, reactive astrocytes are critically important players in the ischemic brain. Careful modulation, rather than inhibition, of reactive gliosis during the right time window therefore appears to be a promising strategy to improve functional recovery after stroke.

## Acknowledgements

The authors acknowledge help with the *in vivo* experiments, especially MRI and behavior tests, by Olivia Käsgen, Veronika Fritz, and Nicole Kuschel. This work was supported by the Friebe Foundation (T0498/28960/16 to MA), Swedish Medical Research Council (2017-02255 to MPy, 2017-00991 to MPa), ALF Gothenburg (724421 to MPy, 716591 to MPa), Söderbergs’ Foundations (to MPy, to MPa), Hjärnfonden (to MPy, to MPa), Hagströmer’s Foundation Millennium (to MPy), Amlöv’s Foundation (to MPy, to MPa), E. Jacobson’s Donation Fund (to UW), the Swedish Stroke Foundation (to MPy, to MPa), EU FP 7 Program TargetBraIn (279017; to MH and MPy), EuroCellNet COST Action (CA15214) including the EuroCellNet COST Action (CA15214) support for Short Scientific Missions (to MA and MH).

## Authors’ contributions

**Conception and design of the study:** MA, UW, MH, MPa, MPy; **acquisition and analysis of data:** MA, UW, FW, AS, YP, FS, NP, LM, MPy.; **drafting a significant portion of the manuscript or figures:** MA, UW, AS, MH, MPa, MPy.

## MATERIAL & METHODS

### Ethics approval

All experiments were conducted in compliance with animal care laws and institutional guidelines and were approved by the Landesamt für Natur, Umwelt und Verbraucherschutz North Rhine-Westphalia, Germany (animal protocol number 84-02.04.2014.A305). This study was done in accordance with the ARRIVE guidelines for reporting *in vivo* animal experiments and the IMPROVE guidelines for stroke animal models (Kilkenny et al., 2010; Percie du Sert et al., 2017).

### Animals and experimental protocol

Animals were housed in individually ventilated cages under 12 h light/12 h darkness cycle with access to water and food *ad libitum*. Two-month-old *GFAP^−/−^Vim^−/−^* (Pekny *et al.*, 1999b) and wild-type (WT) male mice on a C57BL6-129Sv-129Ola mixed genetic background were used (n = 14 WT and n = 15 *GFAP^−/−^Vim^−/−^* mice). Mice with missing values at specific time points (from death during anesthesia or when mice were killed due to post-surgery complications) were excluded (n = 4 WT and n = 7 *GFAP^−/−^Vim^−/−^*). Numbers of mice are presented in the figure legends.

All experiments were planned and recorded in an electronic database to ensure blinded experimentation and restricted access to the data during data recording and analysis (Pallast et al., 2018). The data were analyzed by four raters blinded to the experimental group. The mice underwent photothrombotic stroke and a longitudinal experimental protocol composed of repetitive MRI and three different sensorimotor behavioral tests until 4 weeks after stroke (**Figure 1a**). Photothrombosis was done as described (Aswendt et al., 2021) in the left hemisphere inducing a functional deficit in the right fore- and hindlimb. In brief, after intraperitoneal injection of 1.5 mg of Rose Bengal (in 150 μl of phosphate-buffered saline), 50 mW laser radiation at 561 nm was delivered over 15 min at brain coordinates M/L +2.0 and A/P +0.5 mm, targeting the primary somatosensory forelimb area and primary motor cortex. Post-surgery monitoring included a visual inspection, weighing, and modified neurological deficit scoring (Ito et al., 2018) of general deficits (appearance of eyes and fur, spontaneous movement, epileptic behavior) and focal deficits (whisker response, body/forelimb symmetry, circling behavior, gait). Three behavior tests were used: rotating beam, grid walk and cylinder test. These tests robustly detect sensorimotor deficits as well as recovery for up to 4 weeks post-stroke in mice (Balkaya et al., 2018; Ito *et al.*, 2018; Li et al., 2015; Roome and Vanderluit, 2015). We applied standardized protocols as reported elsewhere (Aswendt *et al.*, 2021). All tests were video recorded and analyzed frame-by-frame by blinded raters (not the experimenters) for hindlimb drops (rotating beam), foot faults (grid walk), and paw drags (cylinder test). An average sensory-motor deficit was calculated from all three tests and normalized to results on poststroke day 3 (P3). Mice were sacrificed by perfusion fixation with phosphate-buffered 4% paraformaldehyde, and the brains were removed for histology.

### MRI data acquisition

MRI data were acquired at the Max Planck Institute for Metabolism Research (Cologne, Germany) using a 94/20USR BioSpec Bruker system (including 660 mT/m B-GA12SHP gradient system, RT-shim and related power supplies, 1H receive-only mouse brain surface coil, and 1H resonator 112/072) and ParaVision v6.01 (Bruker, BioSpin). To reduce movement artefacts and enable reproducible placement of the mouse brain, the mice were anaesthetized initially with isoflurane (2–3% in 70/30 N2/O2) and head-fixed in an animal carrier using tooth and ear bars. Using a custom-made system based on DASYLab (measX), respiration and body temperature were measured (Small Animal Instruments) and electronically recorded, in synchrony with the MRI temporal protocol. Body temperature was maintained at 37°C with a feedback-controlled water circulation system (Medres). After initial adjustments of RF power, shim, and B0 field, a high-resolution, whole-brain, T2-weighted RARE sequence (T2w-MRI) was acquired at coronal slice orientation with a repetition time (TR) of 5500 ms, echo time (TE) of 32.5 ms, flip angle (FA) of 90°, image resolution 0.068×0.068×0.5 mm^3^ (n = 28 slices). Before the start of rs-fMRI, isoflurane anesthesia was continuously reduced to 0.5% after an initial subcutaneous bolus injection of medetomidine (0.1 mg/kg in 0.25 ml of saline) (Domitor, Elanco). Respiration was maintained at 80–120 breaths per minute. A gradient-echo echo planar imaging (GE-EPI) sequence for rs-fMRI was modified from Green et al. (Green *et al.*, 2018) (TR 1420 ms, TE 18 ms, FA 90°, image resolution 0.141×0.141×0.5 mm^3^, and 20 slices). Datasets (raw data and pre-processed data) are publicly available on the German Neuroinformatics Node infrastructure service GIN (https://doi.org/10.12751/g-node.yzjhz3).

### MRI data processing

The raw data in Bruker file format were converted to Nifty format, pre-processed, and registered with the Allen Brain Reference atlas (ccf v3) (Wang et al., 2020) using the in-house developed Python pipeline AIDAmri as described (Pallast et al., 2019). Briefly, the data were converted to Nifty, bias-field corrected, and extracted, followed by a two-step registration. The raw data were untouched, and the transformation was applied to a modified version of the atlas, comprising 98 regions split between hemispheres and selected to match parental regions with the MR image resolution. The AIDAmri code and the modified atlas are available online (https://github.com/aswendtlab/AIDAmri).

Three-dimensional (3D) lesion masks were drawn semi-automatically with ITKsnap (www.itksnap.org) (Yushkevich et al., 2006). The resulting lesion masks at each time point for each group were averaged and presented as incidence maps showing each pixel color-coded to indicate the number of mice in which that pixel was inside the lesion mask. The frequency and fraction of a given brain region that was affected by stroke was also recorded.

The pre-processing of the rs-fMRI data included skull stripping, spatial filtering with a Gaussian kernel sigma of 0.1 mm, slice time correction, motion correction and regression of the respiration, high-pass filtering with a cut-off frequency of 0.01 Hz (100 Hz), and the registration to the defined 98 regions in the standard space of the atlas. The time-series of the regions were calculated as the mean time series of all voxels belonging to the specific region. The correlation matrix was calculated for all 98 regions with Pearson’s correlation on the full dataset. Based on these correlations, graphs were constructed by applying a fixed threshold of 0.1. Functional connectivity data was presented using network science and graph theory in which a brain region relates to a *node* and the connection between two *nodes* to an *edge*. We calculated the universal graph properties *degree* (the number of connections of a node), *edge weight* (connectivity strength between two nodes), *node strength* (the sum of edge weights connected to the node), and *small-worldness* (high local clustering and short average path length between any two nodes). A weighted network analysis taking into account the network edge strength was done with a custom version of the Matlab Brain Connectivity Toolbox (Rubinov and Sporns, 2010), available online (https://github.com/aswendtlab/AIDAconnect).

### Immunohistochemistry

Brains were postfixed overnight in phosphate-buffered 4% paraformaldehyde, dehydrated, embedded in paraffin, and cut into 8-μm sections with a microtome. For immunohistochemistry, sections were rehydrated, subjected to heat-activated antigen retrieval in 0.01 M citrate buffer (pH 6.2, with 2 mM EDTA and 0.05% Tween 20), incubated first with blocking buffer (3% donkey serum and 0.05% Tween 20 in phosphate-buffered saline) and then with primary antibodies diluted in blocking buffer overnight at 4°C. The following primary antibodies were used: rabbit anti-Iba1 (1:400, 019-19741, Wako Chemicals), goat anti-Sox2 (1:200, sc-7379, Santa Cruz Biotechnology), rabbit anti S100 (1:200, Z0311, Dako), mouse anti-Gap43 (1:1000, MAB347, Millipore), and guinea pig anti-Vglut1 (1:500, AB5905, Millipore). Gap43 antibodies were visualized with biotin-conjugated rabbit anti-mouse antibodies (Dako) followed by streptavidin-conjugated Cy3 (Sigma-Aldrich). Iba1, Sox2, S100, and Vglut1 antibodies were visualized with secondary antibodies conjugated with Alexa fluorochromes (Invitrogen).

For assessment of lesion volume, bright-field images of every 24^th^ section stained with hematoxylin and eosin (H&E) were acquired with 2x objective on a Nikon Eclipse 80i light microscope (Bergman Labora). The stroke lesion area was manually delineated with ImageJ (www.imagej.net, NIH).

For assessment of Sox2^+^ and Iba1^+^ astrocytes and microglia, single-plane tiled confocal images covering the motor and somatosensory cerebral cortex of 2 tissue sections 200 μm apart were acquired with a laser-scanning confocal microscope (LSM 700, Carl Zeiss) and standardized acquisition parameters. Square regions of interest (ROIs), 500 μm wide, placed 200 μm from the lesion border and at comparable location in the contralesional cortex, and circular ROIs, 200 μm wide, within the medial and lateral peri-lesion tissue cortical layer 5/6 (as shown in Figure 6a), were assessed for Sox2^+^ cell density and Iba1^+^ area using thresholded images and the Integrated Morphometry Analysis tool in MetaMorph software (Molecular Devices, v. 2.8.5).

For analysis of S100 images, 625-μm-wide square ROIs were placed 100 μm from the lesion border and in contralesional cortex. Directionality was quantified with ImageJ (https://imagej.net/Directionality) and Fourier component to generate histograms with a fitted gaussian curve; amplitude was measured from the baseline of the respective curve. The length of S100^+^ objects was quantified on the same images with the Integrated Morphometry Analysis tool in MetaMorph software. Single-plane 160×160-μm images of Vglut1 and Gap43 immunostained sections were obtained at 1024×1024-pixel resolution by laser scanning confocal microscopy (LSM 700, Zeiss) and standardized acquisition parameters. Images were taken of deep (V-VI) cortical layers in the medial (motor) and lateral (somatosensory) peri-infarct cortex and in the corresponding areas in the contralesional hemisphere. The number, average size, and intensity of positive punctuate structures per image were determined with MetaMorph software.

### Statistics and data visualization

All statistical tests and data plotting were done with Prism (macOS version 8.2.1, www.graphpad.com). Briefly, normality was checked with the D’Agostino-Pearson test or in case of low n with the Shapiro-Wilk test to determine whether to use parametric or nonparametric tests. The parametric tests were t test and one-way and two-way ANOVA, using the sphericity correction method of Geisser and Greenhouse, Sidak’s correction for multiple comparisons between the groups, Dunnett’s test for comparisons of time points, and Tukey’s test for comparison of regions. For the functional connectivity and behavioral analysis, two-way ANOVA with repeated-measures or a mixed-effects model (residual maximum likelihood) was used. The sphericity correction method of Geisser and Greenhouse was applied. For the mixed-effects model, subjects were considered random variables and time and group were considered fixed variables. For functional connectivity analysis only, the original false discovery rate (FDR) correction for multiple comparisons of Benjamin and Hochberg was included. For the results violating the normality tests, we used the non-parametric Mann-Whitney *U* test. Pearson correlation was used to analyze the lesion size determined by MRI, and Spearman correlation was used for the MRI and behavioral data. Immunohistochemistry data were analyzed by one-way ANOVA with Sidak’s or Tukey’s posthoc test. Differences between groups for Iba1, Sox2, and S100 were assessed with Mann-Whitney *U* tests. The data were plotted as box-and-whiskers plot (box extends from the 25^th^ to 75^th^ percentiles with a line in the middle of the box representing the median and 5–95% percentile whiskers), bar graph (mean ± SD), line graph (mean ± SEM), or scatter plot (mean ± SD). Differences were considered significant at p < 0.05.

## Availability of data and materials

The MRI datasets (raw data and pre-processed data) are publicly available in the German Neuroinformatics Node infrastructure repository GIN (https://doi.org/10.12751/g-node.yzjhz3). Other raw and processed data (e.g., microscopy) supporting the conclusions are available upon request. The MRI processing software AIDA is freely available on GitHub (https://github.com/aswendtlab).

