## Supplementary Material for "Reactive Astrocytes Prevent Maladaptive Plasticity after Ischemic Stroke"

### Table 1

Results of Pearson correlation (Pearson r and p-value) of lesion size (MRI P1 and histology P29) and sensorimotor deficit as determined by the number of hindlimb drops (rotating beam test), paw drags (cylinder test), and foot faults (grid walk test) at P3 and P28, respectively.

| | | Lesion size MRI P1 vs. histology P29 | Lesion size MRI P7 vs. histology P29 | Brain tissue loss vs. lesion area |
|---|---|---|---|---|
| ***GFAP$^{-/-}$ Vim$^{-/-}$*** | r | 0.827 | 0.874 | 0.722 |
| | p | 0.014* | 0.010* | 0.043* |
| **WT** | r | 0.845 | 0.712 | 0.928 |
| | p | 0.002** | 0.021* | 0.001*** |
| | | **Lesion size MRI P1 vs. paw drag P3** | **Lesion size MRI P1 vs. hindlimb drop P3** | **Lesion size MRI P1 vs. foot fault P3** |
| | r | 0.397 | 0.073 | 0.137 |
| | p | 0.378 | 0.852 | 0.706 |
| | | **Paw drag P28 vs. lesion size P28** | **Hindlimb drop P28 vs. lesion size P28** | **Foot fault P28 vs. lesion size P28** |
| | r | -0.339 | -0.208 | -0.339 |
| | p | 0.257 | 0.441 | 0.256 |

**Table 2 Top 21 connections from selected regions.**

L - left and R - right hemisphere based on connectivity strength (edge weight).

| | SSp-ul (L - left hemisphere) | | | | SSp-ul (R - right hemisphere) | | | |
|---|---|---|---|---|---|---|---|---|
| | GFAP-/-Vim-/- | | WT | | GFAP-/-Vim-/- | | WT | |
| | *Baseline* | *P28* | *Baseline* | *P28* | *Baseline* | *P28* | *Baseline* | *P28* |
| low | L MB | R SSp-bfd | L LSX | L STRd | L OLF | L HIP | L DORpm | R SSp-ll |
| low | L LSX | R HIP | L SSp-m | L SSp-m | R PTLp | R SSp-n | R mfbs | L HIP |
| low | L SSp-tr | L VIS | R STRd | R HY | R lfbst | L SSp-n | L mfbs | R DORsm |
| low | L SSp-n | R ACA | R cc | R PAL | R SSs | R STRd | L HIP | R HIP |
| low | L HIP | L HIP | L HIP | R cst | R SSp-bfd | R ACA | R SSp-m | R SSp-bfd |
| low | R mfbs | R SSp-ul | R SSp-un | L mfbs | R MOs | R mfbs | R DORpm | R AUD |
| low | R ACA | R cc | L ACA | R ACA | L SSp-ul | R ILA | L ACA | L DORpm |
| middle | R LSX | L STRd | R DORpm | L DORpm | R mfbs | R SSp-bfd | L STRd | L mfbs |
| middle | R DORpm | L MOp | L SSp-bfd | L ACA | R SSp-m | L SSp-ul | L LSX | R SSp-n |
| middle | L MOs | L ACA | R SSp-ll | R mfbs | R HIP | R MOp | L MOp | R DORpm |
| middle | L DORsm | R DORpm | L DORpm | R DORsm | L SSp-ll | L MOp | R SSp-bfd | L LSX |
| middle | L DORpm | L DORpm | L mfbs | R HIP | R cc | R VIS | R SSp-ll | R mfbs |
| middle | L SSp-bfd | R STRd | L STRd | L SSp-bfd | R MB | R DORpm | R cc | R ACA |
| middle | L cc | L SSp-n | R mfbs | L cc | R RSP | R MB | R ACA | R MOp |
| high | L STRd | L mfbs | L SSp-ll | L MOp | R SSp-n | L STRd | R SSp-n | L cc |
| high | L MOp | L SSp-bfd | R ACA | R STRd | L RSP | R cc | L cc | L STRd |
| high | L ACA | L cc | R SSp-ul | R DORpm | R STRd | L SSp-ll | L SSp-ul | R cc |
| high | L mfbs | L SSp-m | L MOp | R cc | R MOp | R RSP | R MOp | R STRd |
| high | L SSp-ll | L SSp-un | L cc | L SSp-ll | R SSp-ll | R SSp-ll | R STRd | L ACA |
| high | L SSp-un | L SSp-ll | L SSp-un | L SSp-un | R SSp-un | R SSp-un | R SSp-un | R SSp-un |

| | SSp-ll (L - left hemisphere) | | | | SSp-ll (R - right hemisphere) | | | |
|---|---|---|---|---|---|---|---|---|
| | GFAP-/-Vim-/- | | WT | | GFAP-/-Vim-/- | | WT | |
| | *Baseline* | *P28* | *Baseline* | *P28* | *Baseline* | *P28* | *Baseline* | *P28* |
| low | R cc | R SSp-ll | R STRd | R SSp-ll | R DORsm | R PTLp | R STRd | L SSp-bfd |
| low | R ACA | R STRd | R cc | L STRd | R RHP | L MB | L SSp-bfd | L RSP |
| low | R STRd | L DORsm | R ACA | R STRd | L SSp-bfd | R STRd | R SSs | L mfbs |
| low | R RSP | L PTLp | L mfbs | R PAL | R MB | L VIS | L HIP | R DORsm |
| low | R mfbs | L ACA | R mfbs | R LSX | L DORpm | L RSP | R cc | R MB |
| low | L ACA | R MOp | L RSP | R DORsm | L STRv | R RHP | L SSp-un | L MB |
| low | L RSP | L mfbs | R MOp | R HIP | R SSs | R DORsm | R SSp-un | R SSp-un |
| middle | L MB | R mfbs | R SSp-ul | L ACA | R MOs | L MOs | L MOp | R mfbs |
| middle | L DORpm | L cc | L SSp-tr | L SSp-un | R cc | L cc | L mfbs | R HIP |
| middle | L cc | R ACA | L HIP | R SSp-bfd | R HIP | R cc | R ACA | R ACA |
| middle | L MOp | L SSp-n | L MOp | R mfbs | R DORpm | R ACA | R SSp-bfd | L SSp-ll |
| middle | L SSp-un | L AUD | R DORpm | L RSP | R RSP | L AUD | R mfbs | L HIP |
| middle | L SSp-bfd | R DORpm | R SSp-un | R ACA | R SSp-n | R mfbs | L ACA | L cc |
| middle | L HIP | L SSp-bfd | L cc | L SSp-bfd | R MOp | L ACA | L DORpm | L DORpm |
| high | R MB | L MOp | R SSp-bfd | R cc | R STRd | L SSp-ll | R MOp | L ACA |
| high | R SSp-ll | L STRd | L DORpm | L DORpm | R SSp-bfd | R MOp | R DORpm | R SSp-bfd |
| high | R DORpm | L DORpm | L SSp-bfd | L cc | L RSP | L SSp-bfd | L cc | L STRd |
| high | R HIP | R SSp-ul | R SSp-ll | R DORpm | L SSp-ll | L MOp | L SSp-ll | R RSP |
| high | L mfbs | R cc | L SSp-un | L MOp | R SSp-un | R SSp-un | L SSp-ul | R DORpm |
| high | L SSp-ul | L SSp-ul | L SSp-ul | L SSp-ul | R SSp-ul | R SSp-ul | R SSp-ul | R SSp-ul |

Table 2 continued.

| | SSp-un (L - left hemisphere) | | | | SSp-un (R - right hemisphere) | | | |
|---|---|---|---|---|---|---|---|---|
| | GFAP-/-Vim-/- | | WT | | GFAP-/-Vim-/- | | WT | |
| | *Baseline* | *P28* | *Baseline* | *P28* | *Baseline* | *P28* | *Baseline* | *P28* |
| low | R cst | L STRd | R SSp-bfd | R SSp-ul | L HIP | R HIP | L SSp-bfd | R HY |
| low | L HIP | R SSp-ul | L MOp | R SSs | R SSs | R MOs | R HY | R LSX |
| low | L PAL | L DORpm | R SSp-ll | R lfbst | L OLF | R MB | R SSp-ll | R AUD |
| low | R ACA | L VIS | R MOp | L ACA | L SSp-tr | L VIS | L HIP | L DORpm |
| low | L SSp-tr | R SSs | R STRd | L mfbs | L SSp-ul | R RHP | L STRd | L HIP |
| low | R DORsm | R STRd | L LSX | R PAL | R DORpm | L STRd | L DORpm | R mfbs |
| low | L ACA | R MB | L SSp-n | L SSp-n | R MB | R cc | R MOp | L MOp |
| middle | R LSX | R SSp-bfd | R SSp-ul | R DORsm | L STRv | L DORpm | R mfbs | L LSX |
| middle | L DORsm | L MB | R ACA | L HIP | L RSP | L SSp-tr | R SSp-n | L cc |
| middle | R mfbs | L SSp-ll | R SSp-un | R STRd | R RSP | R STRd | L SSp-un | R ACA |
| middle | R PAL | R cc | R cc | R HY | R cc | L SSp-n | R cc | R HIP |
| middle | L mfbs | L ACA | L STRd | L SSp-ll | R mfbs | L SSp-bfd | L SSp-ul | L mfbs |
| middle | L MOp | L HIP | R DORpm | R mfbs | R HIP | R RSP | L mfbs | R DORpm |
| middle | L STRd | L cc | L DORpm | R HIP | R RHP | R ILA | L ACA | L STRd |
| high | L SSp-n | L SSp-m | L mfbs | L DORpm | R SSp-n | R SSp-tr | R DORpm | R SSp-n |
| high | L DORpm | L SSs | R mfbs | L cc | R MOp | R ACA | L cc | R SSp-bfd |
| high | L cc | R mfbs | L SSp-ll | R cc | R STRd | R SSp-bfd | R ACA | R cc |
| high | L SSp-ll | L SSp-bfd | L SSp-bfd | R DORpm | R SSp-ll | R SSp-n | R STRd | R STRd |
| high | L SSp-bfd | L SSp-n | L cc | L SSp-bfd | R SSp-bfd | R SSp-ll | R SSp-bfd | L ACA |
| high | L SSp-ul | L SSp-ul | L SSp-ul | L SSp-ul | R SSp-ul | R SSp-ul | R SSp-ul | R SSp-ul |

| | DORsm (L - left hemisphere) | | | | DORsm (R - right hemisphere) | | | |
|---|---|---|---|---|---|---|---|---|
| | | | WT | | | | WT | |
| | *Baseline* | *P28* | *Baseline* | *P28* | *Baseline* | *P28* | *Baseline* | *P28* |
| low | L TEa | L LSX | R SSp-un | R MB | L SSp-ll | R cst | R SSp-un | R ACA |
| low | L mfbs | R MB | R SSp-ul | L AUD | L PTLp | L MOp | R SSp-ul | L ACA |
| low | R SSp-tr | L cst | L MOp | R PAL | R mfbs | L AUD | R AUD | R AUD |
| low | L SSp-un | L AUD | R HIP | L MB | L SSp-bfd | L SSp-bfd | R cst | R PAL |
| low | L PAL | L MB | L ACA | R cc | L RSP | R MB | L SSp-ul | R cst |
| low | L SSp-ll | L MOp | R STRd | L lfbst | L SSp-un | L TEa | R STRd | L SSp-bfd |
| low | L SSp-n | L SSp-bfd | R mfbs | L SSp-bfd | R cc | R mfbs | R mfbs | L STRd |
| middle | L PTLp | L RHP | R HY | R cst | L AUD | R LSX | L HY | R cc |
| middle | L SSp-bfd | L TEa | R SSp-bfd | L STRd | L SSp-ul | L mfbs | R cc | R STRd |
| middle | R HIP | L HY | L lfbst | R STRd | R STRd | L STRd | L mfbs | R lfbst |
| middle | L ACA | L cc | R cc | L cc | R RSP | L HY | R lfbst | R mfbs |
| middle | L STRd | L mfbs | L HY | R mfbs | L ACA | L cc | R HY | L cc |
| middle | L SSp-ul | L STRd | L SSp-bfd | L mfbs | L mfbs | R HY | L SSp-bfd | L mfbs |
| middle | L cc | L lfbst | L STRd | L HY | L cc | R lfbst | R SSp-bfd | L HY |
| high | L AUD | R HY | L cc | R HIP | R lfbst | L MB | L cc | R HY |
| high | L lfbst | R DORsm | L mfbs | R HY | L DORsm | L DORsm | R HIP | R HIP |
| high | L HIP | R HIP | R DORsm | R DORpm | L DORpm | R HIP | L HIP | L HIP |
| high | R DORsm | L HIP | L HIP | R DORsm | L HIP | L HIP | L DORsm | L DORsm |
| high | R DORpm | R DORpm | R DORpm | L HIP | R DORpm | L DORpm | L DORpm | L DORpm |
| high | L DORpm | L DORpm | L DORpm | L DORpm | R HIP | R DORpm | R DORpm | R DORpm |

Table 2 continued.

| | MOp (L - left hemisphere) | | | | MOp (R - right hemisphere) | | | |
|---|---|---|---|---|---|---|---|---|
| | GFAP-/-Vim-/- | | WT | | GFAP-/-Vim-/- | | WT | |
| | *Baseline* | *P28* | *Baseline* | *P28* | *Baseline* | *P28* | *Baseline* | *P28* |
| low | L lfbst | L SSp-ul | L HIP | R DORpm | L STRd | L SSp-bfd | L SSp-un | R SSp-bfd |
| low | L MB | L HIP | L DORsm | R PAL | R DORpm | L SSp-ul | L mfbs | L SSp-bfd |
| low | L SSp-n | R SSp-ll | R SSp-m | R mfbs | R SSp-bfd | L SSs | L DORpm | R LSX |
| low | L DORpm | L SSp-ll | L SSp-un | R SSp-un | L SSp-ul | R AI | R DORpm | R DORpm |
| low | R MB | L mfbs | L SSp-bfd | R MOs | L SSp-ll | L STRd | R SSp-un | R PL |
| low | R STRd | L SSp-bfd | R PL | L mfbs | L RSP | R SSp-ll | R SSp-ll | L LSX |
| low | L SSp-un | R PL | L DORpm | R SSp-ul | R RSP | L SSp-ll | R SSp-n | L ILA |
| middle | L SSp-bfd | R MOs | R SSp-ll | R LSX | R MB | L cc | R SSs | R mfbs |
| middle | R ACA | L STRd | R cc | L SSp-m | L HIP | R RSP | L LSX | R SSp-m |
| middle | L GU | L GU | R STRd | L SSp-ul | L cc | R SSp-ul | L cc | L MOs |
| middle | L ACA | R mfbs | L STRd | R HIP | R SSp-un | L ORB | R cc | L MOp |
| middle | L HIP | R STRd | L mfbs | L cc | R HIP | R cc | L MOs | L cc |
| middle | R LSX | L SSp-m | R MOs | L SSp-ll | R SSp-ll | L VIS | L STRd | L STRd |
| middle | L mfbs | R MOp | R SSp-ul | R MOp | R GU | R ORB | R STRd | R SSp-ul |
| high | L SSp-ll | R cc | R MOp | R STRd | R SSp-m | L PL | L MOp | R STRd |
| high | L cc | L PL | R ACA | L STRd | R cc | R PL | R ACA | R cc |
| high | L SSp-m | L cc | L ACA | R cc | R ACA | L ACA | L ACA | R ILA |
| high | L STRd | L ACA | L SSp-ul | R ACA | R STRd | L MOp | R SSp-ul | L ACA |
| high | L SSp-ul | R ACA | L cc | L ACA | R SSp-ul | R ACA | R MOs | R ACA |
| high | L MOs | L MOs | L MOs | L MOs | R MOs | R MOs | R SSp-m | R MOs |

| | MOs (L - left hemisphere) | | | | MOp (R - right hemisphere) | | | |
|---|---|---|---|---|---|---|---|---|
| | GFAP-/-Vim-/- | | WT | | GFAP-/-Vim-/- | | WT | |
| | *Baseline* | *P28* | *Baseline* | *P28* | *Baseline* | *P28* | *Baseline* | *P28* |
| low | R mfbs | L AUD | L mfbs | R PAL | R RSP | L SSp-un | L LSX | L HIP |
| low | R DORpm | R mfbs | L ILA | L SSp-bfd | L HIP | L AUD | R cc | L SSp-bfd |
| low | L PAL | L VIS | L STRd | R DORpm | L mfbs | R SSp-ul | L PTLp | L ILA |
| low | R HIP | R STRd | L SSp-bfd | L cc | L cc | L RSP | R SSp-ll | L LSX |
| low | R LSX | R GU | L SSp-ll | R SSp-m | L RSP | L SSp-m | L STRd | R SSp-ul |
| low | L SSp-un | L SSs | R SSp-ll | R LSX | R DORsm | L SSp-n | R mfbs | R LSX |
| low | L FRP | L SSp-m | L LSX | L ORB | L RHP | L cc | R SSp-un | L cc |
| middle | L SSp-bfd | R RSP | R PL | R ILA | L PTLp | R STRd | L ORB | R DORpm |
| middle | L SSp-ll | R MOp | L cc | L LSX | R SSp-ul | L SSp-ll | L cc | L STRd |
| middle | R MB | L LSX | R STRd | R cc | L DORpm | R SSp-un | R ORB | R STRd |
| middle | R MOp | R SSp-ll | L SSp-ul | R PL | R DORpm | L VIS | R SSp-ul | R cc |
| middle | R PAL | L STRd | R SSp-m | L STRd | R mfbs | R cc | R SSp-m | L MOp |
| middle | R MOs | R cc | L PL | R SSp-ul | L SSp-ll | R ORB | R STRd | L PL |
| middle | L cc | L cc | L ORB | R STRd | R SSp-ll | L MOs | L SSp-ul | R ILA |
| high | L LSX | R MOs | R SSp-ul | L PL | L MOs | L MOp | L ACA | R ORB |
| high | R ACA | R PL | R ACA | R MOs | R STRd | L PL | R PL | L MOs |
| high | L STRd | R ACA | R MOp | R ACA | L ACA | L ACA | L MOp | R PL |
| high | L SSp-ul | L PL | R MOs | R MOp | R cc | R PL | L MOs | L ACA |
| high | L ACA | L MOp | L ACA | L MOp | R ACA | R MOp | R ACA | R ACA |
| high | L MOp | L ACA | L MOp | L ACA | R MOp | R ACA | R MOp | R MOp |

Table 2 continued.

| | DORpm (L - left hemisphere) | | | | DORpm (R - right hemisphere) | | | |
|---|---|---|---|---|---|---|---|---|
| | GFAP-/-Vim-/- | | WT | | GFAP-/-Vim-/- | | WT | |
| | *Baseline* | *P28* | *Baseline* | *P28* | *Baseline* | *P28* | *Baseline* | *P28* |
| low | L AUD | R cc | R ACA | R ACA | L RSP | L SSp-ll | R ACA | L ACA |
| low | L STRd | L RHP | L lfbst | R AUD | L SSp-tr | R SSp-ul | L SSp-ul | R ACA |
| low | L HY | L SSp-n | R SSp-ul | R SSp-bfd | L STRd | L SSp-ul | L STRd | R AUD |
| low | L PTLp | L SSp-ll | L SSp-ul | L ACA | R cst | L SSp-n | R SSp-un | R PAL |
| low | L SSp-un | R MB | L HY | R PAL | R SSp-ll | R SSp-bfd | L SSp-un | R SSp-bfd |
| low | R cc | L MB | L SSp-un | L SSp-bfd | L cc | L STRd | L SSp-bfd | R STRd |
| low | L SSp-tr | L cst | R STRd | R STRd | R STRd | L HY | L HY | L SSp-bfd |
| middle | L SSp-ll | R STRd | R cc | L STRd | L SSp-ul | L MB | R HY | L STRd |
| middle | L cst | R mfbs | R HY | L HY | L mfbs | L SSp-bfd | R SSp-ul | L HY |
| middle | R mfbs | L HY | R SSp-bfd | R cst | R lfbst | L AUD | R STRd | R cst |
| middle | L SSp-ul | L STRd | R HIP | R cc | L SSp-bfd | L cc | R SSp-bfd | L cc |
| middle | L lfbst | R HIP | R mfbs | R HY | L SSp-ll | L mfbs | L cc | R HY |
| middle | L cc | L cc | L STRd | L cc | R SSp-bfd | R STRd | R cc | R cc |
| middle | L mfbs | L AUD | L SSp-bfd | R mfbs | R cc | R mfbs | L mfbs | L mfbs |
| high | R DORsm | R HY | L cc | R HIP | L DORsm | R HY | R mfbs | L HIP |
| high | R HIP | L mfbs | L mfbs | L mfbs | R DORsm | R HIP | R HIP | R HIP |
| high | L SSp-bfd | R DORsm | R DORsm | L HIP | R HIP | L HIP | L HIP | L DORsm |
| high | L HIP | L DORsm | L HIP | R DORsm | R mfbs | L DORsm | L DORsm | R mfbs |
| high | L DORsm | L HIP | L DORsm | L DORsm | L HIP | R DORsm | R DORsm | R DORsm |
| high | R DORpm | R DORpm | R DORpm | R DORpm | L DORpm | L DORpm | L DORpm | L DORpm |

| | STRd (L - left hemisphere) | | | | STRd (R - right hemisphere) | | | |
|---|---|---|---|---|---|---|---|---|
| | GFAP-/-Vim-/- | | WT | | GFAP-/-Vim-/- | | WT | |
| | *Baseline* | *P28* | *Baseline* | *P28* | *Baseline* | *P28* | *Baseline* | *P28* |
| low | R DORpm | L SSp-ll | L DORsm | L SSp-bfd | R mfbs | L LSX | R SSp-m | L DORsm |
| low | R mfbs | R ACA | R DORpm | R SSp-ul | L STRd | L RHP | L mfbs | R HY |
| low | L SSp-un | R RSP | L SSp-un | R HY | R SSs | L cc | L HY | L SSp-bfd |
| low | R STRd | L MOp | L SSp-ul | R SSp-n | L cc | R ACA | R SSp-un | R SSp-n |
| low | L SSp-ll | R mfbs | L HY | L HIP | R MB | L SSp-ul | R DORpm | R SSp-ul |
| low | L SSp-n | L SSp-bfd | R SSp-ul | R DORsm | L ACA | R RSP | R HY | R DORsm |
| low | L CTXsp | L VIS | R HY | L PAL | R SSp-n | L MB | L DORpm | R SSs |
| middle | L CLA | L RHP | R mfbs | R HIP | R SSp-ll | L VIS | R SSp-n | R HIP |
| middle | L DORsm | L AUD | R MOp | R mfbs | L SSp-ll | R LSX | R MOp | L DORpm |
| middle | R PAL | R HIP | R cc | L DORpm | R HIP | R MB | L PAL | L mfbs |
| middle | L MOs | R cc | L DORpm | L mfbs | L RSP | R mfbs | R mfbs | R DORpm |
| middle | L HIP | L mfbs | R PAL | R DORpm | R DORpm | L AUD | R SSp-ul | R mfbs |
| middle | L LSX | L HIP | L mfbs | R PAL | L HIP | L MOp | R ACA | R ACA |
| middle | R ACA | L HY | R ACA | R LSX | R SSp-un | L HIP | L cc | L cc |
| high | L PAL | L MB | R LSX | L ACA | R RSP | R DORpm | L ACA | L ACA |
| high | R LSX | L DORpm | L ACA | R cc | R ACA | L DORpm | R cc | R cc |
| high | L MOp | R STRd | L PAL | R ACA | R SSp-ul | R HIP | L LSX | R PAL |
| high | L cc | R LSX | L LSX | L cc | R SSp-bfd | L ACA | L STRd | R LSX |
| high | L ACA | L LSX | L cc | L LSX | R cc | L STRd | R LSX | L LSX |
| high | L SSp-ul | L cc | R STRd | R STRd | R MOp | R cc | R PAL | L STRd |

Table 2 continued.

| | SSs (L - left hemisphere) | | | | SSs (R - right hemisphere) | | | |
|---|---|---|---|---|---|---|---|---|
| | GFAP-/-Vim-/- | | WT | | GFAP-/-Vim-/- | | WT | |
| | *Baseline* | *P28* | *Baseline* | *P28* | *Baseline* | *P28* | *Baseline* | *P28* |
| low | L SSp-un | R SSp-ul | R PAL | R ACA | L SSp-ll | L PTLp | R LSX | L HY |
| low | R LSX | R MB | L DORsm | R DORpm | L ORB | R AUD | R ILA | L cc |
| low | R CTXsp | L VIS | L PAL | L mfbs | R DORpm | L STRd | L mfbs | R DORsm |
| low | L SSp-m | L AUD | L MB | L VISC | L RSP | L AUD | R DORpm | L mfbs |
| low | L SSp-tr | R mfbs | R DORsm | R RHP | L eps | R MB | L LSX | R ACA |
| low | R HY | R HY | R SSp-ll | R cc | L STRv | L HIP | L DORpm | L SSp-bfd |
| low | L HIP | L SSp-tr | L VISC | R mfbs | R MOs | R SSp-bfd | L MOp | R HY |
| middle | L DORpm | R DORpm | R SSp-bfd | L DORpm | R PTLp | L MB | L ACA | L HIP |
| middle | L SSp-ul | R ACA | L DORpm | R PAL | R SSp-un | R HIP | R SSp-un | L ACA |
| middle | L mfbs | L HIP | L cst | L SSp-bfd | R HIP | R SSp-un | R SSp-n | R LSX |
| middle | L MOp | R RSP | R SSp-ul | L ACA | R mfbs | R SSp-ul | L cc | R SSp-n |
| middle | L MB | L SSp-un | L SSp-un | R SSp-tr | R cc | R RSP | R cc | L DORpm |
| middle | L ACA | L MB | L SSp-bfd | R STRd | R SSp-ul | L mfbs | L STRd | R PAL |
| middle | R cst | L SSp-m | L SSp-ul | L cc | R SSp-bfd | L SSp-ul | R SSp-bfd | R HIP |
| high | R PAL | L DORpm | L SSp-m | L AUD | R SSp-ll | R LSX | R SSp-ll | L STRd |
| high | L VISC | L cc | R AUD | L HY | R SSp-n | R SSp-n | R SSp-ul | R cc |
| high | L STRd | L STRd | L cc | R HIP | R SSp-m | L SSp-bfd | R STRd | R SSp-bfd |
| high | L cc | R HIP | L STRd | L SSp-n | R RSP | L SSp-un | R ACA | R mfbs |
| high | L SSp-bfd | L SSp-bfd | L SSp-n | L STRd | R STRd | R STRd | R SSp-m | R DORpm |
| high | L SSp-n | L SSp-n | L AUD | L PAL | R VISC | R cc | R MOp | R STRd |
